# Identification of Rab18 as an Essential Host Factor for BKPyV Infection Using a Whole Genome RNA Interference Screen

**DOI:** 10.1101/157602

**Authors:** Linbo Zhao, Michael J. Imperiale

## Abstract

BK polyomavirus (BKPyV) is a human pathogen first isolated in 1971. BKPyV infection is ubiquitous in the human population, with over 80% of adults worldwide being seropositive for BKPyV. BKPyV infection is usually asymptomatic; however, BKPyV reactivation in immunosuppressed transplant patients causes two diseases, polyomavirus-associated nephropathy and hemorrhagic cystitis. To establish a successful infection in its host cells, BKPyV must travel in retrograde transport vesicles to reach the nuclei. To make this happen, BKPyV requires the cooperation of host cell proteins. To further identify host factors associated with BKPyV entry and intracellular trafficking, we performed a whole-genome siRNA screen on BKPyV infection of primary human renal proximal tubular epithelial cells. The results revealed the importance of the Ras-related protein Rab18 and syntaxin 18 for BKPyV infection. Our subsequent experiments implicated additional factors that interact with this pathway, and suggest a more detailed model of the intracellular trafficking process, indicating that BKPyV reaches the ER lumen through a retrograde transport pathway between the late endosome and the ER.

## Introduction

BK polyomavirus (BKPyV) is a small icosahedral DNA virus measuring approximately 45 nm in diameter and was first isolated in 1971 (1). Subsequent serology surveys have revealed that up to 80% of the world’s population has been infected with BKPyV (2), and most of the initial infections occur in early childhood (3). After initial exposure, BKPyV establishes an asymptomatic infection in the urinary tract with periodic shedding into the urine (4). BKPyV reactivation when the immune system is compromised in transplant patients, however, causes two diseases: polyomavirus-associated nephropathy (PVAN) and hemorrhagic cystitis (HC) (5). PVAN is one of the leading causes of graft failure after kidney transplantation (6). Also, BKPyV can be detected in the urine of 90% of allogeneic hematopoietic cell transplant patients who suffer from HC, according to a recent study (7). Even though BKPyV was initially isolated more than 45 years ago, the choices for clinical management of BKPyV reactivation are limited. The first-line treatment for PVAN is lowering the dosage of immunosuppressants, which inevitably increases the risk of acute rejection and graft failure (6). Other options for treating BKPyV reactivation are treatment with cidofovir, leflunomide, and fluoroquinolones; however, the benefit of using these drugs in addition to reducing immunosuppressants has been questioned and has not been investigated carefully in randomized studies (6, 8). Due to the lack of specific antiviral medicines, management of BKPyV reactivation remains a challenge.

BKPyV does not encode any polymerases. As a result, BKPyV exclusively relies on the host DNA replication machinery for its genome replication. To establish a successful infection, BKPyV must deliver its DNA genome into the nucleus to access the host DNA replication machinery, which means crossing the plasma membrane, the ER membrane, and the nuclear envelope (9). Also, BKPyV needs to navigate a crowded cytoplasm to reach the ER. Without any cooperation from host factors, this process would seem difficult if not impossible.

Few host factors have previously been identified that participate in viral entry and intracellular trafficking. To initiate infection, BKPyV binds to gangliosides GD1b or GT1b on the host cell membrane(10). After binding to the cell membrane, BKPyV had been thought to enter host cells via a caveolin-mediated pathway (11), however, subsequent experiments showed that caveolin was dispensable for infecting human renal proximal tubule cells (12). After endocytosis, BKPyV enters the endosome in the same way as other polyomaviruses (13-16). The acidification and maturation of the endosome are essential for BKPyV infection (17), and activates a sorting machinery that involves Rab5, Rab7, Rab9, and Rab11 proteins (14, 15, 18). After sorting through the late endosome, vesicles that contain BKPyV traffic along microtubules and reach the ER at 8-12 hours post infection (17, 19). ER lumen proteins are essential for polyomavirus disassembly and egress from the ER (9, 17, 20-22). Briefly, protein disulfide isomerases induce capsid conformational changes and minor capsid protein exposure (17, 23-28). These exposed minor proteins insert into the ER membrane (29, 30), and partially disassembled polyomaviruses penetrate the ER membrane and enter the cytosol through the endoplasmic reticulum-associated degradation pathway (9, 20, 21, 31, 32). In the cytosol, the nuclear localization signal of the minor capsid proteins guides polyomaviruses into the nucleus via the importin α/β pathway (33-35). In addition to this productive model of entry, a significant portion of internalized polyomavirus enters a non-productive pathway (36). Distinguishing the critical events that BKPyV takes to establish successful infection from other pathways that BKPyV takes remains a challenge.

Numerous assays have been developed for the purpose of identifying host factors associated with viral infections, including genome-wide siRNA screening. By silencing every single human gene with an siRNA pool, and then assessing the effects of the knockdown on infection, it is feasible to dissect the functions of individual host proteins during the viral life cycle. siRNA screening has been extensively applied to research on viruses, such as HIV (37), West Nile Virus (38), HPV (39), VSV (40), and another polyomavirus, SV40 (41). However, no genome-wide siRNA screen has been reported for BKPyV.

Using a whole genome siRNA screen, we have identified a series of potential host factors that are involved in BKPyV infection. DNAJ B14, which has previously been implicated in BKPyV entry, is our top hit, and DNAJ B12 is also among our top 100 hits (21, 41). Most of the other hits we have identified have not been previously reported, however, and many of them are involved in vesicular transport. In this report, we present data showing that two of our primary hits, Rab18 and syntaxin 18, as well as two members of the NRZ complex (RAD50 interactor 1 and ZW10 kinetochore protein), which have previously been shown to interact with Rab18 and syntaxin 18, are essential host factors for BKPyV infection.

## Results

### A high-throughput siRNA screen on BKPyV based on cell cycle analysis

To identify host factors involved in viral entry and intracellular trafficking, we developed and implemented a high-throughput whole-genome siRNA screen for BKPyV infection in its natural host cell, primary human renal proximal tubule epithelial (RPTE) cells. In this screen, BKPyV infection was challenged with the Dharmacon siGENOME Smartpool siRNA library, which includes more than 18,000 human siRNA pools. Each pool contains four unique siRNAs that target the same host gene. The screen was performed in triplicate. Every screen plate included the siGENOME non-targeting control (NTC) as a negative control and a synthetic siRNA which corresponds to the natural BKPyV 5p miRNA targeting the large tumor antigen (TAg) mRNA (siTAg) (42), as a positive control.

Our lab previously showed that BKPyV induces G2/M arrest to take full advantage of the host DNA replication machinery (43); this provided us with a cost-efficient approach to evaluate viral infection in siRNA-transfected cells. By applying a Hoechst DNA stain, cells in G2/M phase are distinguishable from the rest of the cells based on the difference in DNA content in each nucleus. The percentage of G2/M arrested cells (G2%) dramatically increases after BKPyV infection, and we adopted it as the readout for the screening assay.

For the primary screen, RPTE cells were transfected with the siRNA library and cultured for 48 hours to allow depletion of the targeted proteins. Next, cells were infected with BKPyV and cultured for an additional two days. The cells were fixed and stained with Hoechst, and the numbers of nuclei in the different stages of the cell cycle were quantified, recorded, and uploaded to the MScreen web tool developed by the Center for Chemical Genomics (University of Michigan) (44). G2% values were calculated using the MScreen web tool. To proceed with the data analysis, we scaled data from 0 to 100, with 100 representing the effect of our positive control, and 0 representing the effect of negative control. The effect of each siRNA pool was then calculated using this index scale. A larger index value represents a stronger inhibitory effect on the BKPyV infection when an siRNA pool is introduced into RPTE cells, which suggests that the targeted host factor is more likely to be required for BKPyV infection.

Z-factor is a statistical parameter widely used for screen assay evaluation. A minimal Z-factor of 0.5 is required for a reliable siRNA screen (45). The overall Z-factor achieved in our whole primary screen was 0.6, and the R-squared values were approximately 0.9 between replicates. We compared the data from the three replicates in as pairwise manner, as illustrated in the scatter plots in Figure 1. 70% of the siRNA pools in the siGENOME siRNA library inhibited BKPyV infection to some degree when compared to the non-targeting siRNA control. A similar distribution of inhibitory effects was also observed in an siRNA screen using the siGENOME library to assess cellular genes involved in HPV infection (39). The results of the primary screen are in Supplemental Table 1, Sheet1.

**Figure 1.**
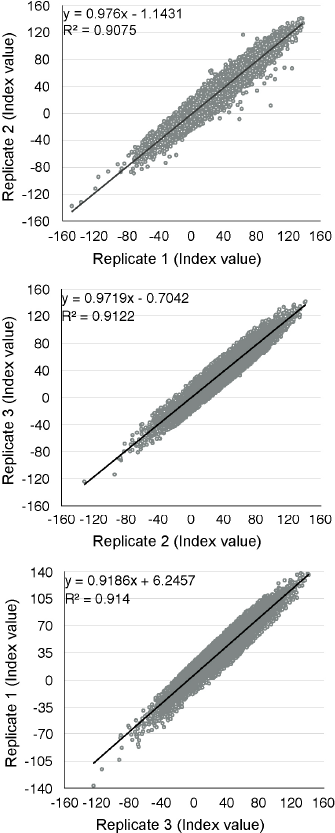
Visualization of whole genome RNAi screen results. The effects of each siRNA pool are normalized to the non-targeting siRNA control (set as 0) and siTAg (set as 100). Index values from the replicates are illustrated in pairwise scatter plots.

We parsed the candidate genes into functional categories and pathways using the Database for Annotation, Visualization, and Integrated Discovery (DAVID) enrichment analysis tool (46, 47). The DAVID analysis showed that translation-related genes were the most enriched group of genes that is required for viral infection among the candidate genes (Supplemental Table 1, Sheet2). Considering that BKPyV exclusively relies on the host protein synthesis machinery, this result is consistent with our expectation. Other than translation-related genes, several clusters of genes that are required for viral infection were also significantly enriched, especially budding and coat proteins associated with vesicular transport. Based on the enrichment analysis results, we manually selected 147 siRNA pools for secondary validation.

G2% is an indirect readout for BKPyV infection. Knocking down cell cycle regulatory proteins may also affect G2/M arrest independent of viral infection; thus false positive or false negative candidates could be introduced into the results during the primary screen. To eliminate these false candidates, we next applied a direct readout, immunofluorescent staining for the early viral protein TAg, to validate these 147 genes. RPTE cells were transfected and infected in 96 well plates with a subset of the siGENOME library that contains the selected 147 siRNA pools. Infected cells were fixed at 48 hours post infection and probed for TAg with primary antibody and FITC-labeled secondary antibody. After acquiring images, the integrated TAg fluorescent intensity per nucleus was recorded, and the inhibition index was calculated as in the primary screen (Supplemental Table 2).

After the secondary screen, DNAJ B14, which has previously been implicated in BKPyV entry, was our top hit, and DNAJ B12 and DNAJC3, which are also involved in polyomavirus trafficking, were also among our top ten hits (21, 41), which indicates the overall robustness of our screen. Rab18 was one of our top validated hits other than these DNAJ proteins. Moreover, syntaxin 18, a protein that has previously been demonstrated to interact with Rab18, also passed our validation.

### The Rab18/NRZ/syntaxin 18 complex is required for BKPyV infection

The Rab protein family is a group of small GTPases that regulate membrane trafficking, and that have been implicated in polyomavirus intracellular trafficking (14, 18, 48). Rab18, however, had not previously been associated with polyomavirus infection. The Rab18 interaction network has been thoroughly investigated (49), and more than 40 proteins have been identified as interacting with Rab18. Among these proteins, syntaxin 18 was also one of our top hits in the screen.

To determine whether Rab18 and syntaxin 18 are required during BKPyV infection, RPTE cells in 12 well plates were transfected with pooled or individual siRNAs targeting these proteins. Cells transfected with the NTC siRNA pool and siTAg served as negative and positive controls, respectively. After transfection, cells were cultured for two days to allow for depletion of the targeted proteins, and then infected with BKPyV at a multiplicity of infection (MOI) of 1 infectious unit (IU)/cell. Cells were lysed at 48 hours post infection, and expression of the targeted proteins, TAg, and β-actin or glyceraldehyde 3-phosphate dehydrogenase (GAPDH) were assessed by Western blotting (Figure 2A). The choice of β-actin or GAPDH was based on avoiding co-migration of the loading control with the protein of interest. Knockdown of Rab18 and syntaxin 18 reduced TAg expression, indicating that both proteins are required for efficient infection. Knockdown of Rab18 entirely blocked BKPyV infection. While there was only partial knockdown of syntaxin 18 with both the pooled or individual siRNAs, TAg expression decreased correspondingly. Because syntaxin 18 is a SNARE protein (50), this suggests that BKPyV enters the ER lumen via a vesicle fusion step.

**Figure 2.**
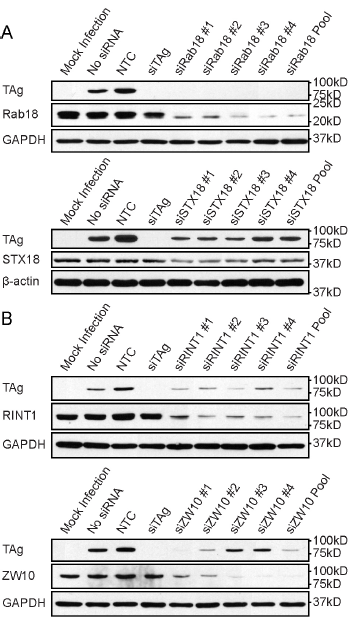
Rab18, syntaxin 18, and the NRZ complex are required for BKPyV infection. RPTE cells were transfected with the indicated siRNAs and then infected with BKPyV. Viral infection (TAg), GAPDH or β-actin expression levels, and knockdown efficiency were examined by Western blot.

In addition to syntaxin 18, Rab18 interacts with ZW10 kinetochore protein (ZW10) and RAD50 Interactor 1 (RINT1), which together with neuroblastoma amplified gene (NAG) are members of the NRZ complex (51). Upon activation by GTP, Rab18 interacts with the ZW10 kinetochore protein from the NRZ complex, thereby forming a Rab18/NRZ/syntaxin 18 complex at the ER (49). In this complex, syntaxin 18 functions as a t-SNARE on the ER membrane where the NRZ components work as a tether to assist with retrograde vesicle docking (51, 52). NRZ/syntaxin 18 cooperate in capturing Rab18-labeled vesicles and initiating the fusion process on the ER membrane. Although none of the NRZ complex proteins were identified in our primary screen, we hypothesized that they are required for BKPyV infection due to their ability to interact with both Rab18 and syntaxin 18. To test whether NRZ components are involved in BKPyV infection, we knocked down RINT1 and ZW10, followed by infection. We could not test NAG due to lack of a useful antibody to measure its expression. The Western blot results indicate that disrupting the NRZ complex by knocking down these two components interferes with infection (Figure 2B).

### Rab18 colocalizes with the viral capsid during BKPyV intracellular trafficking

We next asked whether Rab18 is associated with BKPyV-containing vesicles, since Rab18 targets vesicles to the ER (53, 54). To test this, we fixed RPTE cells at 6 and 8 hours post-infection and stained for the BKPyV major capsid protein, VP1, and Rab18. Images of the stained RPTE cells were acquired with a confocal microscope. The confocal images show that Rab18 and VP1 are partially colocalized at 6 and 8 hours post-infection (Figure 3). This is consistent with the asynchronous trafficking of BKPyV from endosomes after entry (17). These results support our hypothesis that BKPyV travels in Rab18-positive vesicles after being sorted through endosomes.

**Figure 3.**
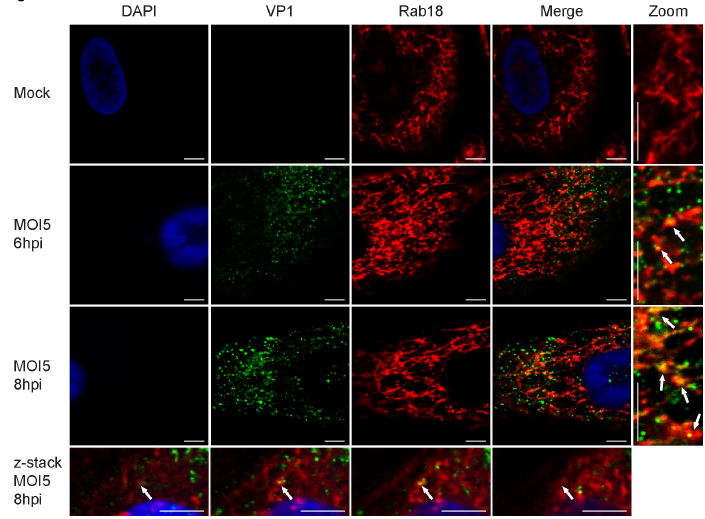
Colocalization of Rab18 and BKPyV capsid protein VP1. RPTE cells were fixed at 6 and 8 hours post infection and were stained for VP1 (green), Rab18 (red), and DAPI (blue). Images were taken using confocal microscopy. Sequential z-stack of images (0.25 μm increments) are illustrated from the bottom of the cell to the top (left to right). White arrows point to VP1-Rab18 colocalization sites. Bars represent 5 μm.

### BKPyV traffics differently after Rab18, syntaxin 18, or NRZ knockdown

To establish a successful infection, BKPyV must reach the ER lumen where disassembly occurs (17). After internalization into host cells, conformational changes of the capsid are neither required nor observed before polyomaviruses enter the ER lumen (27, 28). When polyomavirus particles reach the ER lumen, luminal enzymes disrupt the disulfide bonds between VP1 monomers, thereby initiating disassembly of the viral capsid (26). This process in the ER can be visualized by assaying for VP1 monomer, dimer, and oligomer bands after separation of proteins under non-reducing conditions by SDS-PAGE (17, 28). To confirm that BKPyV requires Rab18 to reach the ER lumen and initiate disassembly, we examined BKPyV disassembly after Rab18 knockdown. RPTE cells were transfected with NTC or Rab18 siRNA for 48 hours and infected with BKPyV at MOI 5. Protein samples were processed and assayed at 24 hours post infection using Western blotting of reducing and non-reducing gels (17, 28). At 0 hours post infection, most of the BKPyV particles were intact and too large to enter the non-reducing SDS-PAGE gel, and no VP1 monomers (42 kDa) or dimers were visible. At 24 hours post infection in cells in which Rab18 was present, BKPyV began to disassemble, and VP1 monomers, dimers, and oligomers could be detected. However, the non-reducing gel suggests that the BKPyV particle could not disassemble efficiently without Rab18 (Figure 4A, top panel). The reducing gel shows that equal amounts of VP1 are detectable in the presence or absence of Rab18, indicating that knocking down Rab18 did not prevent BKPyV from attaching to or entering RPTE cells. We also assessed the effect of knocking down RINT1, ZW10, and syntaxin 18 on BKPyV disassembly (Figure 4A, bottom panel). In each case, lower amounts of VP1 monomers and dimers were detected in the non-reducing gel, suggesting that BKPyV cannot disassemble efficiently without the help of the NRZ complex.

**Figure 4.**
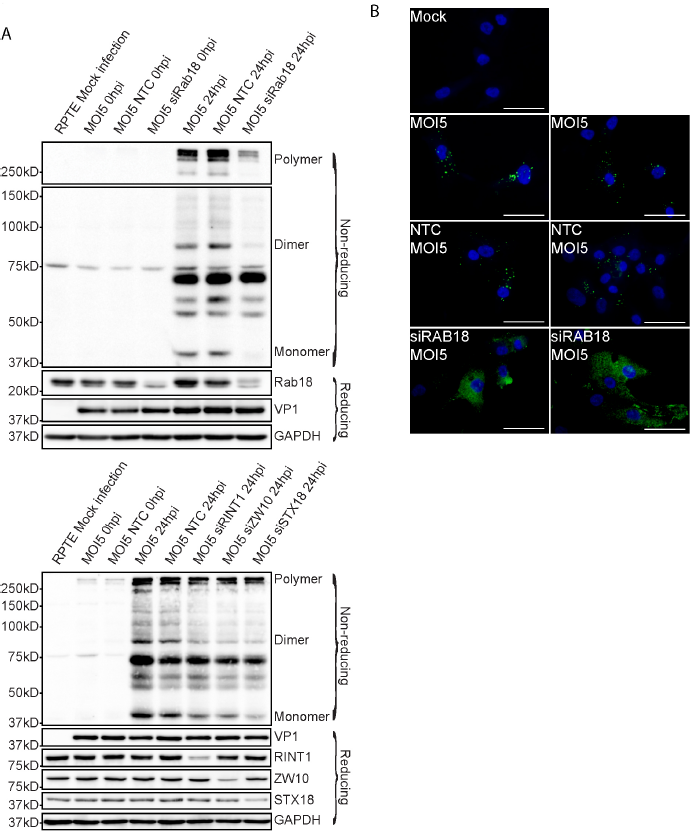
Effects of Rab18 knockdown on BKPyV intracellular trafficking. (A) Rab18, syntaxin 18 (STX18), and NRZ knockdown prevent ER delivery and BKV capsid rearrangement. RPTE cells were transfected with the indicated siRNAs. After 48 hours, they were infected with BKPyV and lysed immediately after adsorption or at 24 hours post-infection under reducing or non-reducing conditions. BKPyV capsid protein VP1, GAPDH, Rab18, STX18, RINT1, and ZW10 levels were examined by Western blot. (B) Alteration of BKPyV intracellular trafficking. RPTE cells were transfected with NTC or siRAB18. BKPyV particles were visualized by immunofluorescent staining for VP1 (green) at 24 hours post infection. Bars represent 100 μm.

Previous studies showed that mouse polyomavirus (MPyV) could enter cells that do not produce proper ganglioside receptors; afterward, these MPyV particles are trapped in a dead-end pathway and cannot establish a successful infection. This indicates that polyomaviruses enter both productive and non-productive pathways inside host cells. These non-productive pathways result in a diffuse distribution of viral particles as detected by immunofluorescent staining (36). In the presence of proper ganglioside receptors, polyomaviruses leave the cell periphery, move along microtubules, and reach the perinuclear area (17, 19, 55, 56). Ultimately, the productive pathway leads to the ER (17, 28, 57). Since we saw that BKPyV could not reach the ER lumen without Rab18 while the entry of BKPyV was not affected (Figure 4A), we wished to address if BKPyV traffics in a different pattern under Rab18 knockdown. To test this, Rab18 was depleted as previously described, and RPTE cells were infected with BKPyV at MOI of 5. Cells were fixed and stained for VP1 at 24 hours post infection (Figure 4B). In the presence of Rab18, BKPyV forms bright foci in the perinuclear area as previously observed (9). However, Rab18 knockdown caused dispersion of the bright perinuclear VP1 foci, and the staining of VP1 appeared to be diffuse and spread throughout the cytoplasm (Figure 4B). Thus, without Rab18 BKPyV traffics in a pattern that is suggestive of a non-productive pathway (36).

### BKPyV is enriched in the late endosome without Rab18

Our previous study demonstrated that acidification of the endosome is essential for BKPyV infection (17), and MPyV colocalizes with the late endosome marker Rab7 after entry (18). Furthermore, BKPyV cannot traffic efficiently to the ER and initiate capsid disassembly without abundant Rab18 and instead appears to be trapped in an unknown compartment(s) diffusely located in the cytoplasm (Figure 4). To identify this compartment, we knocked down Rab18 in RPTE cells, fixed cells at 24 hours post infection, and probed for various organelle markers (Figure 5). We found that the diffuse viral particles do not colocalize with the early endosome marker EEA1, cis-Golgi marker GM130 (58), or trans-Golgi marker Golgin 97 (59). On the other hand, we found that the VP1-rich area overlaps with Rab7. To further confirm this colocalization, images of VP1 and Rab7 were taken using confocal microscopy (Figure 6). We found that the dispersed VP1 partially colocalizes with Rab7 when Rab18 is knocked down, suggesting some viral particles are trapped in the late endosome in the absence of Rab18. There is also some VP1 that does not colocalize with Rab7. This may be a result of asynchronous trafficking of BKPyV, and additional intermediate and dead-end compartments could also exist during the retrograde trafficking step between the endosome and the ER. Our results suggest that BKPyV reaches ER lumen through a Rab18-mediated retrograde transport pathway between the late endosome and the ER.

**Figure 5.**
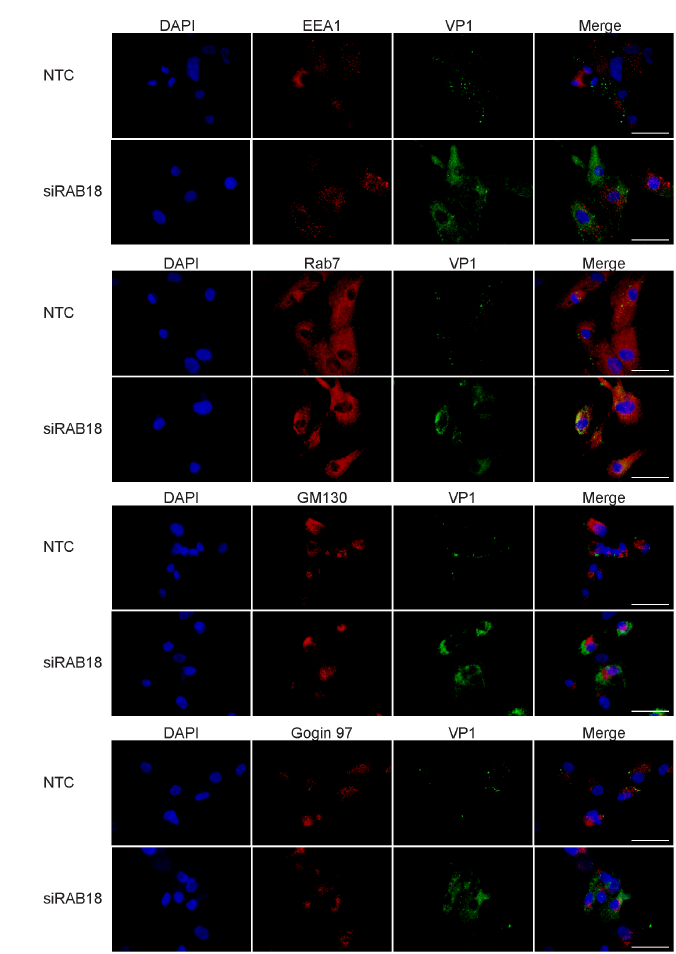
Colocalization of BKPyV and organelle markers. RPTE cells were transfected with NTC or siRAB18, then stained for VP1 (green) and the indicated organelle markers (above each row of panels, red) at 24 hours post infection. Images were taken with an inverted fluorescent microscope. Bars represent 100 μm.

**Figure 6.**
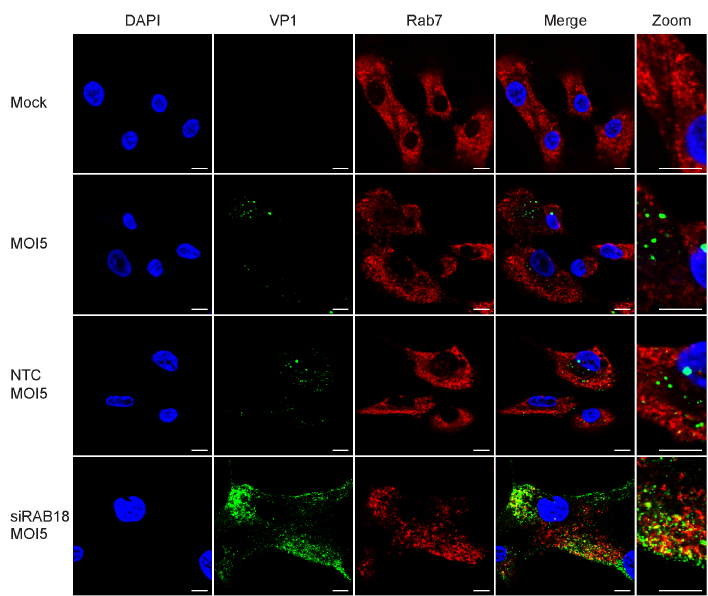
Colocalization of VP1 and Rab7. RPTE cells were transfected with NTC or siRAB18, then stained for VP1 (green) and Rab7 (red) at 24 hours post infection. Images were taken with a confocal microscope. Bars represent 10 μm.

## Discussion

BK polyomavirus (BKPyV) was initially isolated more than 45 years ago (1), however, our understanding of the early events in the BKPyV life cycle contains gaps. Polyomaviruses appear to take advantage of multiple endocytic pathways to enter the host cell. However, some of the pathways that BKPyV uses are non-productive (36), which suggests that identifying host factors solely by morphological observations, especially protein co-localization evidence, may be deceiving. SV40, BKPyV, MPyV, and cholera toxin have been observed to enter the cell via a caveolin-mediated pathway (11, 60-64). However, subsequent experiments have demonstrated that caveolin is dispensable (12, 14, 65-68). Because of the limitations of the available biological tools, research of the earliest stages of the polyomavirus life cycle has progressed relatively slowly.

Whole genome siRNA screening provides an unbiased tool to investigate some of the remaining questions. However, one of the biggest challenges of developing an siRNA screen assay is finding a readout that is sensitive, cost efficient, and suitable for high throughput automation. One of the most common readouts for a viral siRNA screen is cell viability. However, BKPyV, along with many viruses, does not lyse host cells within a reasonable period compared to the duration of the RNAi effect in cells. This behavior of BKPyV makes cell viability measurements impractical. Another common strategy to evaluate viral infection is by incorporating reporter genes into the viral genome. Because BKPyV is highly sensitive to genomic modification, several approaches attempting to integrate reporter genes into the BKPyV genome have failed (data not shown). Therefore, constructing modified viral particles containing a reporter gene was not practical for us. Finally, immunofluorescent staining for viral protein expression is neither cost- nor time-efficient for processing hundreds of plates. For all these reasons, we needed a different readout for evaluating viral infection.

The percentage of cells in G2/M phase provided us a cost-efficient assay for the primary screen. Our primary screen results indicated that knocking down 70% of the human genes in RPTE cells inhibited BKPyV infection to some degree when compared to the non-targeting control. A similar data distribution was also observed in an siRNA screen for HPV infection using the same siRNA library (39). We found that the NTC siRNA reproducibly slightly increased BKPyV infection compared to the no siRNA control. This could be because of interference of exogenous siRNA with the host intrinsic miRNA processing and silencing mechanism. BKPyV encodes a miRNA that downregulates TAg mRNA levels (42, 69). If the introduction of exogenous siRNA affects the miRNA processing and silencing mechanism, TAg expression would increase due to lower levels of the mature BKPyV miRNA.

The increased BKPyV infection induced by the non-targeting siRNA control could also be an off-target effect of RNAi. While validating our primary screening results, we found many of our primary hits were not reproducible when tested with individual siRNAs instead of siRNA pools. It appears that off-target effects of RNAi are inevitable for whole genome siRNA screens, and an ‘ideal non-targeting’ siRNA pool does not exist (70). The most likely cause of the off-target effect is the passenger strand of the siRNA. Each siRNA carries a complimentary strand (passenger strand) to enhance the stability of the siRNA molecules. This passenger strand can also be integrated into the RNA-induced silencing complex (RISC), which possibly increases off-target effects. Moreover, even if only the guide strand enters the RISC complex, RISC prefers the 2nd to the 8th nucleotides of that strand, called the seed region, to recognize targeted mRNAs (70-74). Considering this characteristic of RNAi, each seven-nucleotide seed region can target more than one gene. Off-target effects can therefore increase the expense of validation and lower the efficiency of screening.

After validating some of the top hits as well as other genes that are known to be involved in the same vesicular trafficking pathway, Rab18, syntaxin 18, ZW10, and RINT1 were confirmed as essential host factors for BKPyV infection. Rab18 has been associated with lipid droplet homeostasis and vesicular transport between the Golgi apparatus and the ER (53, 54). The majority of Rab18 usually localizes to the membrane of the ER and the cis-Golgi apparatus (53). Also, Rab18 is recruited to the surface of lipid droplets, the endosome, and the lysosome (75, 76). Rab18 has been shown to interact with syntaxin 18, ZW10, and RINT1 by affinity chromatography followed by mass spectrometry (49). Based on previous studies on yeast proteins, ZW10 and RINT1 interact with each other via their N-termini and interact with the NAG protein to form the NRZ complex. The t-SNARE protein syntaxin 18 on the ER membrane indirectly attaches to the NRZ complex (51, 52, 77). The overall structure of the NRZ complex is a protrusion about 20 nm in length from the surface of the ER membrane (78). GTP-activated Rab18 on the surface of vesicles interacts with ZW10, thereby tethering vesicles to the ER membrane. Syntaxin 18 is a SNARE protein and mediates vesicle fusion to the ER (50). After the NRZ complex captures vesicles, syntaxin 18 mediates vesicle fusion with the ER membrane.

We propose a model of BKPyV entry and intracellular trafficking to the ER based on previous findings and our current results (Figure 7). To initiate infection, BKPyV first binds to its ganglioside receptors GT1b and GD1b (10). After initial attachment, BKPyV is believed to form deep invaginations of the host plasma membrane in the same manner as SV40 (79). The length of ceramide tails of the ganglioside plays a critical role in formation of the invagination (79). Next, BKPyV enters host cells via a ganglioside-dependent, caveolin- and clathrin-independent pathway (12, 80). Meanwhile, glycoproteins on the plasma membrane appear to serve as traps for polyomavirus (81): viral particles that bind to glycoproteins will eventually enter non-productive pathways. Only those particles that bind to the correct ganglioside receptor can enter the productive infection pathway (36). Furthermore, binding to the gangliosides is sufficient to drive transport of artificial particles to the ER (18). This indicates that binding to the proper ganglioside is both necessary and sufficient for BKPyV to traffic in retrograde transport vesicles to the ER. In addition, conformational changes of the capsid are neither required nor observed before polyomaviruses enter the ER lumen (27, 28). These findings reveal that BKPyV may remain bound to the membrane throughout the intracellular trafficking until reaching the ER lumen. BKPyV enters the endosome in the same manner as other polyomaviruses (13-15). Acidification of the endosome activates a ganglioside-sorting machinery that sorts BKPyV into secondary vesicles along with gangliosides (17, 18, 82). Without Rab18, we found that BKPyV accumulates in the late endosome (marked by Rab7). This suggests that Rab18 mediates late endosome to ER trafficking of BKPyV and that it buds and traffics together with virus-containing vesicles along microtubules (17, 19, 55, 83). After the NRZ complex on the ER membrane captures and tethers vesicles to the ER surface, syntaxin 18 further interacts with a yet to be identified v-SNARE on the vesicles, and the syntaxin 18/v-SNARE complex mediates vesicle fusion to the ER membrane. This allows successful entry of BKPyV into the ER lumen. Our finding supports the conclusion that late endosome to ER trafficking plays a critical role in BKPyV infection.

**Figure 7.**
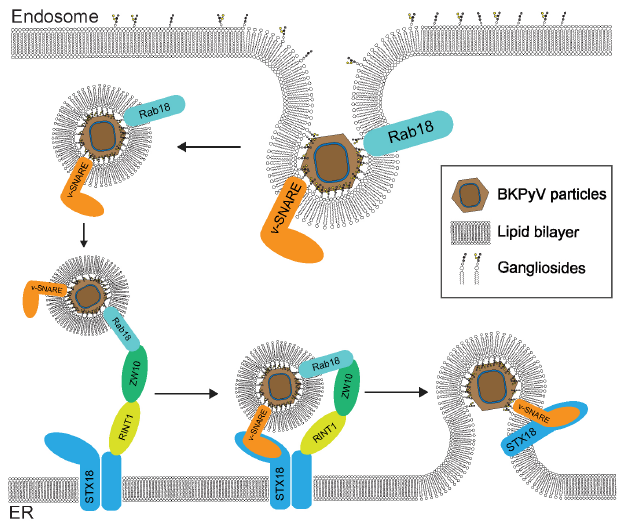
Model of BKPyV vesicular trafficking. (A) BKPyV enters a vesicle from the membrane of the Golgi apparatus or the endosome. (B) GTP-bound Rab18 interacts with ZW10 of the NRZ tethering complex. (C) Syntaxin 18 on the ER membrane interacts with v-SNARE. (D) Syntaxin 18 and v-SNARE mediate vesicle fusion.

It has been shown that fluorescently labeled gangliosides can spontaneously transport in a retrograde direction to the ER (77). BKPyV and the other polyomaviruses are therefore like hitchhikers that take a ride along this ganglioside-mediated retrograde pathway to the ER. Whether additional interactions between the viral capsid and host proteins play a role in BKPyV trafficking to the ER is still unclear. Gangliosides play important roles in both endocytosis and retrograde trafficking. Polyomaviruses and several toxins are proposed to take advantage of lipid-mediated endocytosis and retrograde trafficking pathways (reviewed by Ewers and Helenius (80)). Beside polyomaviruses, some other non-enveloped viruses infect cells in a similar manner; therefore, they may also utilize part of this lipid-mediated retrograde trafficking pathway to establish an infection (84): The retrograde paths of both HPV and BKPyV can be blocked with retrograde traffic inhibitors retro-1, retro-2, and BFA (17, 39, 85, 86). Norovirus also binds to gangliosides (87, 88), and forms deep invaginations on model membranes (89). Gangliosides have been demonstrated to be important for rotavirus infection (90), and rotavirus also traffics to and is sorted through the endosome (91). However, details of the Rab18-mediated pathway are largely unknown, and further efforts will be needed to fully reveal the details of this retrograde trafficking pathway.

## Acknowledgements

We would like to thank the members of the Imperiale Lab, past and present, for their support and encouragement; Adam Lauring, Akira Ono, and Billy Tsai for critical review of the manuscript and sharing reagents with us; and everyone, especially Martha Larsen and Nick Santoro, from the University of Michigan Center for Chemical Genomics for their efforts in implementing the siRNA screen. Research reported in this publication was supported in part by the National Cancer Institute of the National Institutes of Health (NIH) under award number P30CA046592 to the University of Michigan Comprehensive Cancer Center (UMCCC), funding from the Cancer Research Committee of the UMCCC, NIH grant AI060584 awarded to M.J.I., and a Rackham Graduate Student Research Grant awarded to L.Z.

## Materials and Methods

Cell culture. Primary renal proximal tubule epithelial (RPTE) cells purchased from Lonza were maintained in the recommended medium, REGM BulletKit (REGM/REBM, Lonza, CC-3190), at 37°C with 5% CO2 in a humidified incubator. Cells recovered from a single frozen vial from Lonza (Passage 2) were cultured for three passages. Afterward, cells at passage 5 were passaged one more time and plated for the screening, or aliquoted and frozen in liquid nitrogen for later experiments. For all experiments other than the primary screen, frozen aliquots (Passage 5) were recovered about one week before each experiment, and cells were then plated for experiments.

Infection. BKPyV (Dunlop) was cultured, purified on a cesium chloride linear gradient, and titered as described previously (17, 92). RPTE cells were infected as follows at two days post siRNA transfection. Cells were pre-chilled for 15 min at 4°C. Purified viruses were diluted to 175,000 IU/ml (MOI 1) or 875,000 IU/ml (MOI 5) in REBM/REGM. 400 μl of the diluted virus were added to the wells of a 12 well plate and incubated at 4°C for 1 hour with shaking every 15 minutes to distribute the inoculum over the entire well. The plate was transferred to 37°C after the 1-hour incubation.

siRNA and siRNA library. The whole-genome human siGENOME smart pool siRNA library from Dharmacon was acquired and prepared by the Center for Chemical Genomics (University of Michigan). All other siRNAs were also purchased from Dharmacon: Non-targeting siRNA control (D-001206-14); Rab18 siRNA (LQ-010824-00); STX18 siRNA (LQ-020624-01); ZW10 siRNA (LQ-003948-00); RINT1 siRNA (LQ-004976-01); siRNA targeting large T antigen (custom synthesized with the sequence 5’ AUCUGAGACUUGGGAAGAGCAU 3’), which corresponds to the natural BKPyV 5p miRNA (42). Primary siRNA screening in 384 well plates. The siRNA library was rehydrated at 500 nM in siRNA buffer (Dharmacon, B-002000-UB-100) according to the Basic siRNA Resuspension protocol from Dharmacon. 1 μl 500nM siRNA suspension was spotted into each well of 384-well PE Viewplates on a Biomek laboratory automation workstation. RPTE cells were transfected according to the Lipofectamine RNAiMAX (Thermo Fisher Scientific) manual. Briefly, transfection complexes were prepared by adding 9 μl of diluted transfection reagent (0.78% RNAiMAX reagent v/v in REBM/REGM without antibiotics) to each well of the 384-well plates onto which the siRNAs had been spotted. The transfection complexes were incubated at room temperature for 20 min before adding 1,800 cells suspended in 10 μl REBM/REGM without antibiotics to each well. Transfected cells were cultured at 37°C for 48 h, after which cells were infected by the following procedure: incubate plates at 4°C for 15 min; dilute purified BKPyV Dunlop in cold REGM/REBM; dispense 5 μl 1,800,000 IU/ml virus to each well with a Multidrop Combi reagent dispenser (Thermo Fisher Scientific); incubate plates at 4°C for 1 hour; transfer plates to 37°C for additional 48 hours. Next, cells were fixed with 4% paraformaldehyde (Electron Microscopy Sciences) at room temperature for 20 min, permeabilized with 0.1% Triton x-100 (MilliporeSigma) in PBS for 5 min at room temperature, and stained with 2 μg/ml Hoechst 33342 (Thermo Fisher Scientific, H3570) in PBS for 15 min at room temperature. Cells were washed three times with PBS after each staining step. Images of the wells were taken with an ImageXpress Micro XLS high-throughput microscope and analyzed with MetaXpress High-Content Image Acquisition and Analysis software. The quantified data generated from MetaXpress were uploaded and further analyzed with MScreen, a high-throughput analysis system developed by the Center for Chemical Genomics (University of Michigan) (44). Low-quality data points generated from wells with contamination or significant cell death were discarded.

Secondary siRNA screening in 96 well plate. For siRNA transfection in 96 well plates, the primary screening protocol was scaled up by applying 3 times the volume used in 384 wells plate. In addition to the Hoechst stain, cells were also incubated for 1 hour each with 5% goat serum, anti-TAg monoclonal antibody (pAb416) at 1:200 dilution in 5% goat serum (93), and goat anti-mouse IgG–FITC (MilliporeSigma, F2012) at 1:200 dilution in 5% goat serum. Cells were rinsed with 1x PBS 3 times between each step. Images of the wells were taken with an ImageXpress Micro XLS high-throughput microscope and analyzed with MetaXpress High-Content Image Acquisition and Analysis software for TAg intensity.

Tertiary validation in 12 well plates. siRNAs were rehydrated at 1 μM in siRNA buffer (Dharmacon, B-002000-UB-100). Transfection complexes were prepared by mixing 20 μl of 1 μM siRNA with 380 μl of diluted transfection reagent (0.74% RNAiMAX reagent v/v in REBM/REGM without antibiotics) in each well of a 12-well plate. The transfection complexes were incubated at room temperature for 20 min before adding 70,000 cells suspended in 400 μl REBM/REGM without antibiotics to each well. RPTE cells were infected at two days post transfection.

Preparation of protein lysates. Cells were lysed at 48 hours post infection with E1A buffer [50 mM HEPES (pH 7), 250 mM NaCl, and 0.1% NP-40, with inhibitors: 5 μg/ml PMSF, 5 μg/ml aprotinin, 5 μg/ml leupeptin, 50 mM sodium fluoride and 0.2 mM sodium orthovanadate added right before use] (MilliporeSigma). For non-reducing gels, cells were rinsed with 1x PBS with 10 mM N-Ethylmaleimide (MilliporeSigma, E3876) and harvested by scraping. Harvested cells were further pelleted with a centrifuge and lysed with a Triton lysis buffer [10 mM Tris (pH 7.6), 10 mM sodium phosphate, 130 mM NaCl, 1% Triton X-100, 10 mM NEM, protease inhibitors (Roche, 11697498001)]. Insoluble cell debris was removed by centrifuging lysed cells at 16,100 xg and discarding the pellet. Protein concentration was quantified with the Bradford assay (Bio-Rad).

Western blotting. Protein samples were separated on 12% SDS-PAGE gels. After electrophoresis, the proteins were transferred to a nitrocellulose membrane (MilliporeSigma, pore size 0.2 μm) in Towbin transfer buffer (25 mM Tris, 192 mM glycine, 20% methanol) at 60 V overnight. Membranes were blocked with 2% nonfat milk in PBS-T buffer (144 mg/L KH_2_PO_4_, 9 g/L NaCl, 795 mg/L Na_2_HPO_4_, pH 7.4, 0. 1% Tween 20) for 1 hour. Membranes were probed with primary and secondary antibodies diluted in 2% milk in PBS-T as follows: TAg (pAb416) at 1:5,000 dilution (93); syntaxin 18 (Abcam, ab156017) diluted in 5% BSA at 1:1,000; Rab18 (MilliporeSigma, SAB4200173) at 1:10,000; VP1(pAb5G6) 1:1000; GAPDH (Abcam, ab9484) at 1: 10,000; ZW10 kinetochore protein (Abcam, ab53676) at 1:300; RAD50 interactor 1 (MilliporeSigma, HPA019875) at 1:200; β-actin (Cell signaling, #4967) at 1:10,000; horseradish peroxidase (HRP)-conjugated ECL sheep anti-mouse (GE healthcare, NA931V) at 1: 5,000; and HRP-conjugated ECL donkey anti-rabbit antibody (GE Healthcare, NA934V) at 1: 5,000. Protein bands were visualized with HRP substrate (Millipore, WBLUF0100) and exposure to X-ray film or the Syngene PXi gel doc system.

Immunofluorescent staining. 18 mm circular coverslips (#1.5 thickness, Electron Microscopy Sciences, 72222) were coated with 0.1% poly-L-lysine in water (MilliporeSigma, P8920) in 12 well plates for 5 min at room temperature. Coverslips were then rinsed with cell culture grade water and allowed to dry for at least 2 hours. RPTE cells were then seeded onto the coverslips in the 12-well plate. For processing, the cells were fixed with 4% PFA at room temperature for 20 min. Antigen was retrieved with antigen retrieval buffer (100 mM Tris, 5% [w/v] urea, pH 9.5) at 95 °C for 10 min. Cell membranes were permeabilized with 0.1% Triton X-100 in PBS at room temperature for 5 min. Coverslips were blocked with 5% goat serum in PBS for 1 hour and probed with diluted primary and secondary antibody successively, 1 hour each. Coverslips were washed with PBS for three times between each step. Lastly, coverslips were mounted with Prolong Gold Reagent with DAPI (Thermo Fisher Scientific, P36931). Primary or secondary antibodies were diluted in 5% goat serum (MilliporeSigma) as follows: VP1 (pAb5G6) at 1:200; Rab18 (MilliporeSigma, SAB4200173) at 1:1500; Rab7 (Cell Signaling, #9367) at 1:100; GM130 (Cell Signaling, #12480) at 1:200; EEA1 (Cell Signaling, #3288) at 1:200; Golgin-97 (Cell Signaling, #13192) at 1:100; goat anti-mouse IgG–FITC (MilliporeSigma, F2012) at 1:200; goat anti-rabbit IgG-DL594 (Thermo Fisher Scientific, 35561) at 1:200. Images were taken with Olympus BX41 or Leica inverted SP5 confocal microscope system with a 100x objective. Confocal images were acquired and processed with LAS AF, LAS X software from Leica.

## References

1. Gardner SD, Field AM, Coleman DV, Hulme B. 1971. New human papovavirus (B.K.) isolated from urine after renal transplantation. The Lancet 1:1253–1257.

2. Egli A, Infanti L, Dumoulin A, Buser A, Samaridis J, Stebler C, Gosert R, Hirsch HH. 2009. Prevalence of Polyomavirus BK and JC Infection and Replication in 400 Healthy Blood Donors. J Infect Dis 199:837–846.

3. Kean JM, Rao S, Wang M, Garcea RL. 2009. Seroepidemiology of human polyomaviruses. PLoS Pathog 5:e1000363.

4. Ahsan N, Shah KV. 2006. Polyomaviruses and human diseases. Adv Exp Med Biol 577:1–18.

5. Jiang M, Abend JR, Johnson SF, Imperiale MJ. 2009. The role of polyomaviruses in human disease. Virology 384:266–273.

6. Kuypers DRJ. 2012. Management of polyomavirus-associated nephropathy in renal transplant recipients. Nat Rev Nephrol 8:390–402.

7. Lunde LE, Dasaraju S, Cao Q, Cohn CS, Reding M, Bejanyan N, Trottier B, Rogosheske J, Brunstein C, Warlick E, Young JAH, Weisdorf DJ, Ustun C. 2015. Hemorrhagic cystitis after allogeneic hematopoietic cell transplantation: risk factors, graft source and survival. Bone Marrow Transplant 50:1432–1437.

8. Ramos E, Drachenberg CB, Wali R, Hirsch HH. 2009. The Decade of Polyomavirus BK-Associated Nephropathy: State of Affairs. Transplantation 87:621–630.

9. Bennett SM, Jiang M, Imperiale MJ. 2013. Role of cell-type-specific endoplasmic reticulum-associated degradation in polyomavirus trafficking. J Virol 87:8843–8852.

10. Low JA, Magnuson B, Tsai B, Imperiale MJ. 2006. Identification of gangliosides GD1b and GT1b as receptors for BK virus. J Virol 80:1361–1366.

11. Moriyama T, Marquez JP, Wakatsuki T, Sorokin A. 2007. Caveolar endocytosis is critical for BK virus infection of human renal proximal tubular epithelial cells. J Virol 81:8552–8562.

12. Zhao L, Marciano AT, Rivet CR, Imperiale MJ. 2016. Caveolin- and clathrin-independent entry of BKPyV into primary human proximal tubule epithelial cells. Virology 492:66–72.

13. Querbes W, O’Hara BA, Williams G, Atwood WJ. 2006. Invasion of host cells by JC virus identifies a novel role for caveolae in endosomal sorting of noncaveolar ligands. J Virol 80:9402–9413.

14. Liebl D, Difato F, Horníková L, Mannová P, Štokrová J, Forstová J. 2006. Mouse polyomavirus enters early endosomes, requires their acidic pH for productive infection, and meets transferrin cargo in Rab11-positive endosomes. J Virol 80:4610–4622.

15. Engel S, Heger T, Mancini R, Herzog F, Kartenbeck J, Hayer A, Helenius A. 2011. Role of endosomes in simian virus 40 entry and infection. J Virol 85:4198–4211.

16. Ashok A, Atwood WJ. 2003. Contrasting roles of endosomal pH and the cytoskeleton in infection of human glial cells by JC virus and simian virus 40. J Virol 77:1347–1356.

17. Jiang M, Abend JR, Tsai B, Imperiale MJ. 2009. Early events during BK virus entry and disassembly. J Virol 83:1350–1358.

18. Qian M, Cai D, Verhey KJ, Tsai B. 2009. A lipid receptor sorts polyomavirus from the endolysosome to the endoplasmic reticulum to cause infection. PLoS Pathog 5:e1000465.

19. Eash S, Atwood WJ. 2005. Involvement of cytoskeletal components in BK virus infectious entry. J Virol 79:11734–11741.

20. Inoue T, Tsai B. 2015. A nucleotide exchange factor promotes endoplasmic reticulum-to-cytosol membrane penetration of the nonenveloped virus simian virus 40. J Virol 89:4069–4079.

21. Bagchi P, Walczak CP, Tsai B. 2015. The Endoplasmic Reticulum Membrane J Protein C18 Executes a Distinct Role in Promoting Simian Virus 40 Membrane Penetration. J Virol 89:4058–4068.

22. Chromy LR, Oltman A, Estes PA, Garcea RL. 2006. Chaperone-mediated in vitro disassembly of polyoma-and papillomaviruses. J Virol 80:5086–5091.

23. Norkin LC, Anderson HA, Wolfrom SA, Oppenheim A. 2002. Caveolar Endocytosis of Simian Virus 40 Is Followed by Brefeldin A-Sensitive Transport to the Endoplasmic Reticulum, Where the Virus Disassembles. J Virol 76:5156–5166.

24. Rainey-Barger EK, Mkrtchian S, Tsai B. 2007. Dimerization of ERp29, a PDI-like protein, is essential for its diverse functions. Mol Biol Cell 18:1253–1260.

25. Schelhaas M, Malmström J, Pelkmans L, Haugstetter J, Ellgaard L, Grünewald K, Helenius A. 2007. Simian Virus 40 depends on ER protein folding and quality control factors for entry into host cells. Cell 131:516–529.

26. Walczak CP, Tsai B. 2011. A PDI family network acts distinctly and coordinately with ERp29 to facilitate polyomavirus infection. J Virol 85:2386–2396.

27. Nelson CDS, Ströh LJ, Gee GV, O’Hara BA, Stehle T, Atwood WJ. 2015. Modulation of a pore in the capsid of JC polyomavirus reduces infectivity and prevents exposure of the minor capsid proteins. J Virol 89:3910–3921.

28. Inoue T, Dosey A, Herbstman JF, Ravindran MS, Skiniotis G, Tsai B. 2015. ERdj5 Reductase Cooperates with Protein Disulfide Isomerase To Promote Simian Virus 40 Endoplasmic Reticulum Membrane Translocation. J Virol 89:8897–8908.

29. Daniels R, Rusan NM, Wadsworth P, Hebert DN. 2006. SV40 VP2 and VP3 insertion into ER membranes is controlled by the capsid protein VP1: implications for DNA translocation out of the ER. Molecular Cell 24:955–966.

30. Rainey-Barger EK, Magnuson B, Tsai B. 2007. A chaperone-activated nonenveloped virus perforates the physiologically relevant endoplasmic reticulum membrane. J Virol 81:12996–13004.

31. Inoue T, Tsai B. 2011. A large and intact viral particle penetrates the endoplasmic reticulum membrane to reach the cytosol. PLoS Pathog 7:e1002037.

32. Geiger R, Andritschke D, Friebe S, Herzog F, Luisoni S, Heger T, Helenius A. 2011. BAP31 and BiP are essential for dislocation of SV40 from the endoplasmic reticulum to the cytosol. Nature Cell Biology 13:1305–1314.

33. Nakanishi A, Shum D, Morioka H, Otsuka E, Kasamatsu H. 2002. Interaction of the Vp3 nuclear localization signal with the importin α2/β heterodimer directs nuclear entry of infecting simian virus 40. J Virol 76:9368–9377.

34. Nakanishi A, Itoh N, Li PP, Handa H, Liddington RC, Kasamatsu H. 2007. Minor capsid proteins of simian virus 40 are dispensable for nucleocapsid assembly and cell entry but are required for nuclear entry of the viral genome. J Virol 81:3778–3785.

35. Bennett SM, Zhao L, Bosard C, Imperiale MJ. 2015. Role of a nuclear localization signal on the minor capsid proteins VP2 and VP3 in BKPyV nuclear entry. Virology 474:110–116.

36. You J, O’Hara SD, Velupillai P, Castle S, Levery S, Garcea RL, Benjamin T. 2015. Ganglioside and Non-ganglioside Mediated Host Responses to the Mouse Polyomavirus. PLoS Pathog 11:e1005175.

37. Zhou H, Xu M, Huang Q, Gates AT, Zhang XD, Castle JC, Stec E, Ferrer M, Strulovici B, Hazuda DJ, Espeseth AS. 2008. Genome-scale RNAi screen for host factors required for HIV replication. Cell Host Microbe 4:495–504.

38. Krishnan MN, Ng A, Sukumaran B, Gilfoy FD, Uchil PD, Sultana H, Brass AL, Adametz R, Tsui M, Qian F, Montgomery RR, Lev S, Mason PW, Koski RA, Elledge SJ, Xavier RJ, Agaisse H, Fikrig E. 2008. RNA interference screen for human genes associated with West Nile virus infection. Nature 455:242–245.

39. Lipovsky A, Popa A, Pimienta G, Wyler M, Bhan A, Kuruvilla L, Guie M-A, Poffenberger AC, Nelson CDS, Atwood WJ, DiMaio D. 2013. Genome-wide siRNA screen identifies the retromer as a cellular entry factor for human papillomavirus. Proc Natl Acad Sci USA 110:7452–7457.

40. Lee AS-Y, Burdeinick-Kerr R, Whelan SPJ. 2014. A genome-wide small interfering RNA screen identifies host factors required for vesicular stomatitis virus infection. J Virol 88:8355–8360.

41. Goodwin EC, Lipovsky A, Inoue T, Magaldi TG, Edwards APB,Van Goor KEY, Paton AW, Paton JC, Atwood WJ, Tsai B, DiMaio D. 2011. BiP and multiple DNAJ molecular chaperones in the endoplasmic reticulum are required for efficient simian virus 40 infection. MBio 2:e00101–11.

42. Seo GJ, Fink LHL, O’Hara B, Atwood WJ, Sullivan CS. 2008. Evolutionarily conserved function of a viral microRNA. J Virol 82:9823–9828.

43. Jiang M, Zhao L, Gamez M, Imperiale MJ. 2012. Roles of ATM and ATR-Mediated DNA Damage Responses during Lytic BK Polyomavirus Infection. PLoS Pathog 8:e1002898.

44. Jacob RT, Larsen MJ, Larsen SD, Kirchhoff PD, Sherman DH, Neubig RR. 2012. MScreen. J Biomol Screen 17:1080–1087.

45. Zhang J, Chung T, Oldenburg K. 1999. A Simple Statistical Parameter for Use in Evaluation and Validation of High Throughput Screening Assays. J Biomol Screen 4:67–73.

46. Da Wei Huang, Sherman BT, Lempicki RA. 2009. Systematic and integrative analysis of large gene lists using DAVID bioinformatics resources. Nat Protoc 4:44–57.

47. Da Wei Huang, Sherman BT, Lempicki RA. 2008. Bioinformatics enrichment tools: paths toward the comprehensive functional analysis of large gene lists. Nucl Acids Res 37:1–13.

48. Mannová P, Forstová J. 2003. Mouse polyomavirus utilizes recycling endosomes for a traffic pathway independent of COPI vesicle transport. J Virol 77:1672–1681.

49. Gillingham AK, Sinka R, Torres IL, Lilley KS, Munro S. 2014. Toward a comprehensive map of the effectors of rab GTPases. Dev Cell 31:358–373.

50. Iinuma T, Aoki T, Arasaki K, Hirose H, Yamamoto A, Samata R, Hauri H-P, Arimitsu N, Tagaya M, Tani K. 2009. Role of syntaxin 18 in the organization of endoplasmic reticulum subdomains. J Cell Sci 122:1680–1690.

51. Tagaya M, Arasaki K, Inoue H, Kimura H. 2014. Moonlighting functions of the NRZ (mammalian Dsl1) complex. Front Cell Dev Biol 2:25.

52. Hirose H, Arasaki K, Dohmae N, Takio K, Hatsuzawa K, Nagahama M, Tani K, Yamamoto A, Tohyama M, Tagaya M. 2004. Implication of ZW10 in membrane trafficking between the endoplasmic reticulum and Golgi. EMBO J 23:1267–1278.

53. Dejgaard SY, Murshid A, Erman A, Kizilay O, Verbich D, Lodge R, Dejgaard K, Ly-Hartig TBN, Pepperkok R, Simpson JC, Presley JF. 2008. Rab18 and Rab43 have key roles in ER-Golgi trafficking. J Cell Sci 121:2768–2781.

54. Liu S, Storrie B. 2012. Are Rab proteins the link between Golgi organization and membrane trafficking? Cell Mol Life Sci 69:4093–4106.

55. Zila V, Difato F, Klimova L, Huerfano S, Forstová J. 2014. Involvement of microtubular network and its motors in productive endocytic trafficking of mouse polyomavirus. PLoS ONE 9:e96922.

56. Sanjuan N, Porrás A, Otero J. 2003. Microtubule-dependent intracellular transport of murine polyomavirus. Virology 313:105–116.

57. Magnuson B, Rainey EK, Benjamin T, Baryshev M, Mkrtchian S, Tsai B. 2005. ERp29 triggers a conformational change in polyomavirus to stimulate membrane binding. Molecular Cell 20:289–300.

58. Nakamura N. 1995. Characterization of a cis-Golgi matrix protein, GM130. J Cell Biol 131:1715–1726.

59. Follit JA. 2006. The Intraflagellar Transport Protein IFT20 Is Associated with the Golgi Complex and Is Required for Cilia Assembly. Mol Biol Cell 17:3781–3792.

60. Parton RG. 1994. Regulated internalization of caveolae. J Cell Biol 127:1199–1215.

61. Orlandi PA, Fishman PH. 1998. Filipin-dependent Inhibition of Cholera Toxin: Evidence for Toxin Internalization and Activation through Caveolae-like Domains. J Cell Biol 141:905–915.

62. Richterová Z, Liebl D, Horák M, Palková Z, Stokrová J, Hozák P, Korb J, Forstová J. 2001. Caveolae are involved in the trafficking of mouse polyomavirus virions and artificial VP1 pseudocapsids toward cell nuclei. J Virol 75:10880–10891.

63. Pelkmans L, Kartenbeck J, Helenius A. 2001. Caveolar endocytosis of simian virus 40 reveals a new two-step vesicular-transport pathway to the ER. Nature Cell Biology 3:473–483.

64. Eash S, Querbes W, Atwood WJ. 2004. Infection of vero cells by BK virus is dependent on caveolae. J Virol 78:11583–11590.

65. Damm E-M, Pelkmans L, Kartenbeck J, Mezzacasa A, Kurzchalia T, Helenius A. 2005. Clathrin- and caveolin-1-independent endocytosis: entry of simian virus 40 into cells devoid of caveolae. J Cell Biol 168:477–488.

66. Gilbert JM, Goldberg IG, Benjamin TL. 2003. Cell Penetration and Trafficking of Polyomavirus. J Virol 77:2615–2622.

67. Shogomori H, Futerman AH. 2001. Cholera toxin is found in detergent-insoluble rafts/domains at the cell surface of hippocampal neurons but is internalized via a raft-independent mechanism. J Biol Chem 276:9182–9188.

68. Torgersen ML, Skretting G, van Deurs B, Sandvig K. 2001. Internalization of cholera toxin by different endocytic mechanisms. J Cell Sci 114:3737–3747.

69. Broekema NM, Imperiale MJ. 2013. miRNA regulation of BK polyomavirus replication during early infection. In.

70. Tschuch C, Schulz A, Pscherer A, Werft W, Benner A, Hotz-Wagenblatt A, Barrionuevo LS, Lichter P, Mertens D. 2008. Off-target effects of siRNA specific for GFP. BMC Mol Biol 9:60.

71. Jackson AL, Bartz SR, Schelter J, Kobayashi SV, Burchard J, Mao M, Li B, Cavet G, Linsley PS. 2003. Expression profiling reveals off-target gene regulation by RNAi. Nat Biotechnol 21:635–637.

72. Jackson AL, Burchard J, Schelter J, Chau BN, Cleary M, Lim L, Linsley PS. 2006. Widespread siRNA “off-target” transcript silencing mediated by seed region sequence complementarity. RNA 12:1179–1187.

73. Birmingham A, Anderson EM, Reynolds A, Ilsley-Tyree D, Leake D, Fedorov Y, Baskerville S, Maksimova E, Robinson K, Karpilow J, Marshall WS, Khvorova A. 2006. 3’ UTR seed matches, but not overall identity, are associated with RNAi off-targets. Nat Meth 3:199–204.

74. Mohr S, Bakal C, Perrimon N. 2010. Genomic screening with RNAi: results and challenges. Annual review of biochemistry 79:37–64.

75. Zhang J, Chang D, Yang Y, Zhang X, Tao W, Jiang L, Liang X, Tsai H, Huang L, Mei L. 2017. Systematic investigation on the intracellular trafficking network of polymeric nanoparticles. Nanoscale 9:3269–3282.

76. Zhang J, Zhang X, Liu G, Chang D, Liang X, Zhu X, Tao W, Mei L. 2016. Intracellular Trafficking Network of Protein Nanocapsules: Endocytosis, Exocytosis and Autophagy. Theranostics 6:2099–2113.

77. Civril F, Wehenkel A, Giorgi FM, Santaguida S, Di Fonzo A, Grigorean G, Ciccarelli FD, Musacchio A. 2010. Structural analysis of the RZZ complex reveals common ancestry with multisubunit vesicle tethering machinery. Structure 18:616–626.

78. Ren Y, Yip CK, Tripathi A, Huie D, Jeffrey PD, Walz T, Hughson FM. 2009. A structure-based mechanism for vesicle capture by the multisubunit tethering complex Dsl1. Cell 139:1119–1129.

79. Ewers H, Römer W, Smith AE, Bacia K, Dmitrieff S, Chai W, Mancini R, Kartenbeck J, Chambon V, Berland L, Oppenheim A, Schwarzmann G, Feizi T, Schwille P, Sens P, Helenius A, Johannes L. 2010. GM1 structure determines SV40-induced membrane invagination and infection. Nature Cell Biology 12:11-8-sup pp 1–12.

80. Ewers H, Helenius A. 2011. Lipid-mediated endocytosis. Cold Spring Harb Perspect Biol 3:a004721–a004721.

81. Qian M, Tsai B. 2010. Lipids and proteins act in opposing manners to regulate polyomavirus infection. J Virol 84:9840–9852.

82. Chinnapen DJ-F, Hsieh W-T, Welscher te YM, Saslowsky DE, Kaoutzani L, Brandsma E, D’Auria L, Park H, Wagner JS, Drake KR, Kang M, Benjamin T, Ullman MD, Costello CE, Kenworthy AK, Baumgart T, Massol RH, Lencer WI. 2012. Lipid sorting by ceramide structure from plasma membrane to ER for the cholera toxin receptor ganglioside GM1. Dev Cell 23:573–586.

83. Moriyama T, Sorokin A. 2008. Intracellular trafficking pathway of BK Virus in human renal proximal tubular epithelial cells. Virology 371:336–349.

84. Taube S, Jiang M, Wobus CE. 2010. Glycosphingolipids as receptors for non-enveloped viruses. Viruses 2:1011–1049.

85. Nelson CDS, Carney DW, Derdowski A, Lipovsky A, Gee GV, O’Hara B, Williard P, DiMaio D, Sello JK, Atwood WJ. 2013. A retrograde trafficking inhibitor of ricin and Shiga-like toxins inhibits infection of cells by human and monkey polyomaviruses. MBio 4:e00729–13.

86. Carney DW, Nelson CDS, Ferris BD, Stevens JP, Lipovsky A, Kazakov T, DiMaio D, Atwood WJ, Sello JK. 2014. Structural optimization of a retrograde trafficking inhibitor that protects cells from infections by human polyoma- and papillomaviruses. Bioorg Med Chem 22:4836–4847.

87. Taube S, Perry JW, Yetming K, Patel SP, Auble H, Shu L, Nawar HF, Lee CH, Connell TD, Shayman JA, Wobus CE. 2009. Ganglioside-linked terminal sialic acid moieties on murine macrophages function as attachment receptors for murine noroviruses. J Virol 83:4092–4101.

88. Han L, Tan M, Xia M, Kitova EN, Jiang X, Klassen JS. 2014. Gangliosides are ligands for human noroviruses. J Am Chem Soc 136:12631–12637.

89. Rydell GE, Svensson L, Larson G, Johannes L, Römer W. 2013. Human GII.4 norovirus VLP induces membrane invaginations on giant unilamellar vesicles containing secretor gene dependent α1,2 fucosylated glycosphingolipids. Biochimica et Biophysica Acta (BBA) - Biomembranes 1828:1840–1845.

90. Martínez MA, López S, Arias CF, Isa P. 2013. Gangliosides have a functional role during rotavirus cell entry. J Virol 87:1115–1122.

91. Arias CF, Silva-Ayala D, López S. 2015. Rotavirus entry: a deep journey into the cell with several exits. J Virol 89:890–893.

92. Abend JR, Low JA, Imperiale MJ. 2007. Inhibitory effect of gamma interferon on BK virus gene expression and replication. J Virol 81:272–279.

93. Harlow E, Whyte P, Franza BR, Schley C. 1986. Association of adenovirus early-region 1A proteins with cellular polypeptides. Mol Cell Biol 6:1579–1589.

